# Actin architecture steers microtubules in active cytoskeletal composite

**DOI:** 10.1101/2022.08.03.502658

**Authors:** Ondřej Kučera, Jérémie Gaillard, Christophe Guérin, Manuel Théry, Laurent Blanchoin

## Abstract

Cytoskeletal motility assays use surface-immobilised molecular motors to propel cytoskeletal filaments. These assays have been widely employed to characterise the motor properties and interactions of cytoskeletal elements with themselves or with external factors. Moreover, the motility assays are a promising class of bio-inspired active tools for nanotechnological applications. While effective utilisation of these assays involves controlling the filament direction and speed, either as a sensory readout or a functional feature, designing a subtle control embedded in the assay is an ongoing challenge. Here we investigate the interaction between motor-propelled microtubules and networks of actin filaments. We demonstrate that the microtubules respond to a network of actin filaments and that this response depends on the network’s architecture. Both linear actin filaments and a network of actin branched by the Arp2/3 complex decelerate microtubule gliding; however, an unbranched actin network provides additional guidance and effectively steers the microtubules. This effect, which resembles the recognition of cortical actin architecture by microtubules, is a conceptually new means of controlling the filament gliding in the motility assay with potential application in the design of active materials and cytoskeletal nano-devices.

## Introduction

Microtubules and actin filaments are biopolymers that form the cytoskeleton in eukaryotic cells. Through their dissipative self-assembly, crosstalk, interaction with motor proteins and a plethora of regulatory factors, these cytoskeletal structures underlie fundamental biological processes such as transport, motility, and morphogenesis (*1, 2*). The building blocks of the cytoskeletal machinery can be, thanks to their self-organising properties, isolated from cells and reconstructed into functional assemblies *in vitro*. Transcending the original purpose of the reconstitution biology, i.e. reducing the complexity of biological machinery under controlled conditions, the *in vitro* cytoskeletal experimentation has led to the development of active materials (*3*) and nano-devices for sensory applications or non-classical computing (*4–7*). A great portion of artificial cytoskeletal systems makes use of gliding assay. In this assay, cytoskeletal filaments are propelled along a surface by ATP consuming motor proteins that have been adsorbed to this very surface. An effective application of the gliding assay involves regulation of the filament gliding, either as a sensory readout or a functional feature. It can be achieved by the modulation of the motor activity (*8–10*) or by steering the gliding direction by geometrical constraints (*11–16*) and external forces (*17–20*).

An interesting opportunity to regulate the filament gliding stems from the interaction of filaments with themselves—the underpinning of collective phenomena (*3*)—by the assistance of molecular linkers (*21–26*) or depletion forces (*27, 28*). Although the main focus has been on single species of biopolymers, either microtubules or actin filaments, recent works inspired by cytoskeletal crosstalk in cells (*2*) have shown that combining the cytoskeletal components in a composite brings additional qualities to gliding assays (*29–31*). Nevertheless, the main qualitative property of cytoskeletal crosstalk, its dependence on the actin architecture (*2*), remained unexplored.

Here, we scrutinise the interaction landscape between motile microtubules and the two types of actin architecture: linear actin filaments or a network of actin branched by the Arp2/3 complex (*32*). To this end, we developed a microtubule gliding assay in the presence of a layer of actin filaments. In this system, we observed that actin filaments influence the microenvironment experienced by a microtubule by imposing friction and providing guidance cues by alignment. Notably, the microtubule gliding was directionally controlled by actin architecture and its anisotropic structural geometry. This effect, which is reminiscent of recognising cortical actin architecture by microtubules in cells (*2, 32*), can be exploited to design active materials and cytoskeletal nano-devices.

## Results

We attached kinesin-I motors to a glass surface at the density of 17.4 μm^-2^ and let them propel pre-polymerised GMPCPP microtubules in the presence of linear actin architecture, or a network branched using Arp2/3 complex (Fig. 1 a, b, Methods). Microtubules were introduced at a low concentration reducing the probability of their collisions during gliding. The untethered actin mesh was polymerised *in situ* with the aid of a crowding agent (Fig. 1 c). Using TIRF microscopy imaging, we monitored the growth of the actin and waited until the actin fluorescence intensity reached a plateau when we started the observation of gliding microtubules (Fig. 1 d). We note that actin mesh covered the entire field of view at this time, forming a relatively homogeneous “carpet” with a distinctive internal organisation given by the actin architecture (Fig. 1 e).

**Figure 1.**
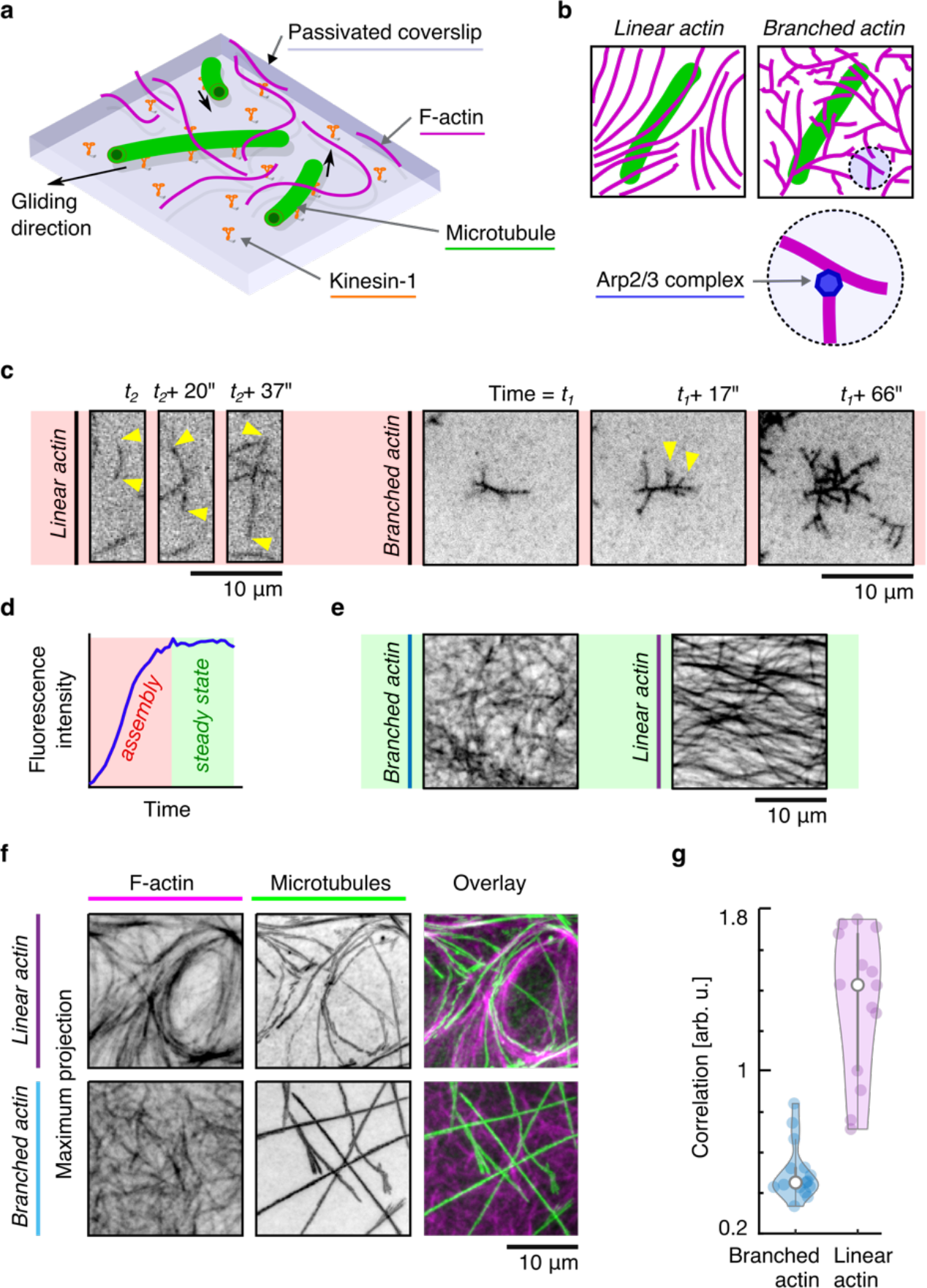
Microtubule gliding assay in the presence of actin networks. **a**, Schematic representation of the assay. **b**, Schematic representation of a microtubule in linear and branched actin network. **c**, Time-lapse intensity inverted fluorescence micrographs of polymerisation of linear actin and branched actin filaments. The yellow arrowheads indicate the ends of the filament (linear actin) and newly grown branches (branched actin). **d**, The temporal profile of actin fluorescence intensity reveals the polymerisation state of actin filaments. **e**, Intensity inverted fluorescence micrographs of branched and linear actin networks in the saturation condition. **f**, Maximum projection images of actin networks and microtubules reveal the microtubule guidance by linear actin. **g**, Correlation of maximum projections of corresponding actin and microtubule images (*N* = v = 14 and 15 per condition, *n* = 3). Individual data points are accompanied by violin plots: Central marks represent the median, top and bottom edges of the box indicate the 75th and 25th percentiles, respectively. Whiskers extend the 95% confidence intervals.

We observed that microtubules visibly followed the local organisation in the linear actin mesh as if the actin served as a template for microtubule trajectories (Fig. 1 f). The measurement of the angle between microtubules and the dominant local orientation of actin filaments confirmed that microtubules tend to align with linear actin (median: 8°, 75% percentile: 18°, maximum angle: 89°, N = 380, v = 10, n = 3). This effect, which resembles cortical microtubule guidance, was, by contrast, unnoticeable in the branched actin network (Fig. 1 f), which lacks geometric features that could provide guidance. The correlation of maximum temporal projection of microtubule trajectories (projection time 180 s) with the actin geometry (Fig. 1 g) confirmed the difference as highly reproducible (Mann-Whitney U-test of equal medians, p = 9.43 · 10^-6^). Since we did not observe transport or visible deformation of actin by motile microtubules, we presume that this actin-architecture-specific behaviour of motile microtubules stems from a geometry-related difference in the actin microenvironment.

Compared to control in the absence of actin, we observed that the gliding of microtubules in both actin architectures appeared slower (Fig. 2 a), indicating that the actin network acts as a mechanical resistance. Occasionally, in the presence of actin filaments, microtubules exhibited markedly complex velocity profiles with pausing or quick switching of the gliding direction, leading sporadically to a quasi-oscillatory movement (Fig. S1). These observations indicate inhomogeneities, non-linear effects, the presence of elastic coupling between the polymers, or a complex combination of these factors which will be difficult to fit into a more quantitative picture.

**Figure 2.**
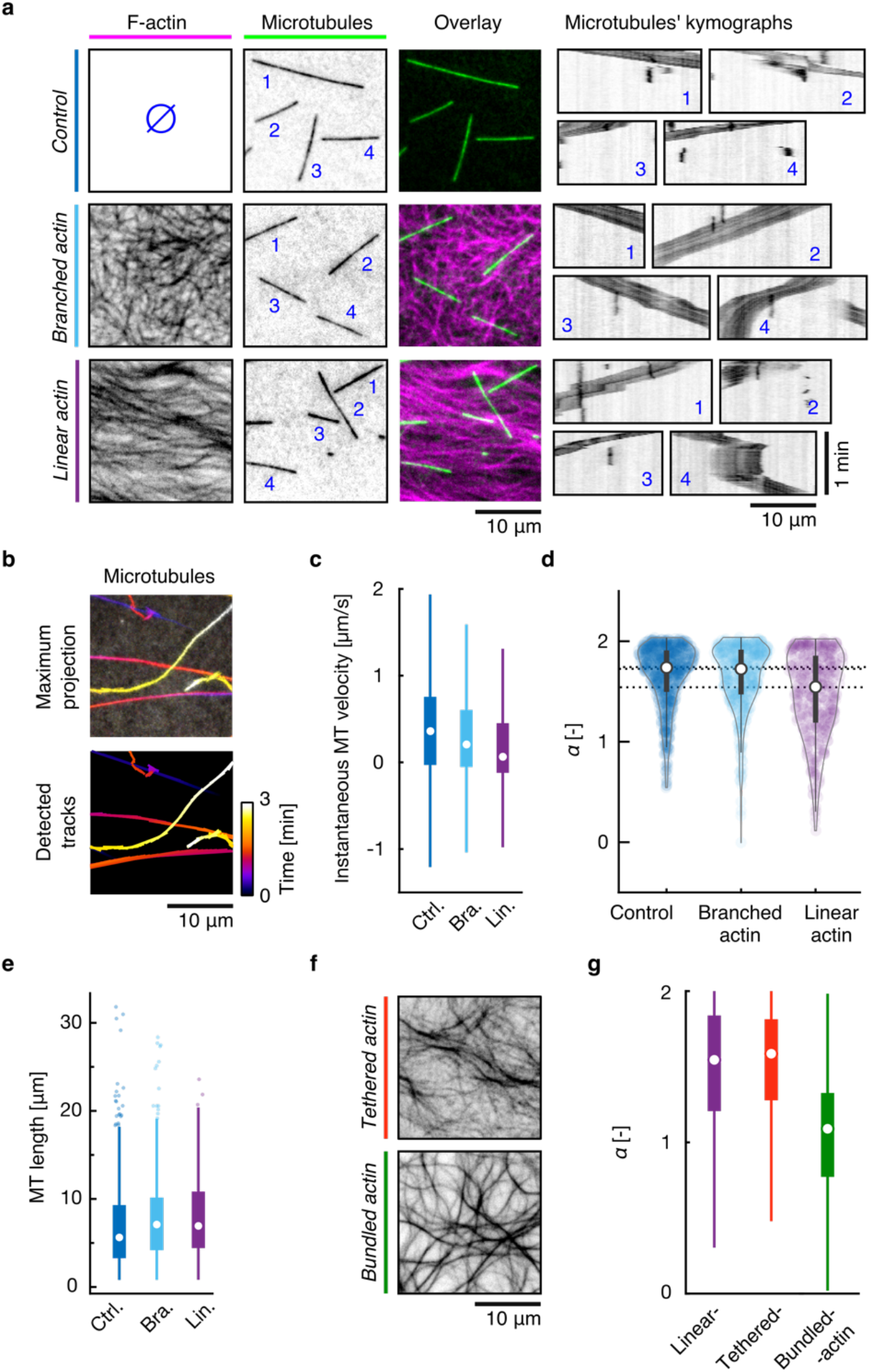
Microtubule motility is controlled by actin architecture. **a**, Typical fluorescence micrographs of actin and microtubules with corresponding kymographs of microtubule motility. **b**, An example of temporal colour-coded maximum projection of microtubule mobility and corresponding detected microtubule tracks. **c**, Box plot of instantaneous microtubule velocities (outliers not shown). **d**, Diffusion exponent, α, for all detected microtubule tracks (Control: *N* = 666, v = 9, *n* = 3. Branched actin: *N* = 479, v = 10, *n* = 3. Linear actin: *N* = 380, v = 10, *n* = 3). **e**, Comparison of detected microtubule lengths in control experiments and in the presence of branched and linear actin. **f**, Inverse intensity fluorescence micrographs of tethered and bundled linear actin filaments. **g**, The comparison of the diffusion exponent of microtubules gliding in the three types of linear actin (Linear actin: *N* = 380, v = 10, *n* = 3. Tethered actin: *N* = 2452, v = 7, *n* = 2. Bundled actin: *N* = 114, v = 21, *n* = 2.). Central marks represent the median, top and bottom edges of the box indicate the 75th and 25th percentiles, respectively. Whiskers extend the 95% confidence intervals.

We tracked all microtubules in our assay to quantify the microtubule response to their microenvironment. The information about microtubule position at each time-point of the observation gave us access to microtubule trajectories (Fig. 2 b) and instantaneous velocities (Fig. 2 c). Since very short trajectories cannot be reliably analysed, in the following, we selected only those microtubules that covered a distance of at least 6 μm, which is an experimentally tuned compromise between the spatial-temporal resolution of the image acquisition and the data fitting.

First, using the trajectories, we plot the mean-squared displacement, MSD, of the microtubules (Fig. S2 a). MSD is coupled with the delay, or the time interval over which it is measured, by a power law, MSD ∝ τ^α^. The value of the diffusion exponent, a, demarcates three regimes of the motion: sub-diffusion related to confinement (0 < α < 1), random diffusion (α ≈ 1), and super-diffusion (1 < α <2). By curve fitting, we determined the values of the diffusion exponents from our experimental data (R^2^ = 0.98, N = 1525 microtubule trajectories). The comparison of the distributions of the diffusion exponent (Fig. 2 d) revealed that the values measured in all groups predominantly correspond to super-diffusive motion (1 < α < 2), in concordance with the visual inspection of the microtubules’ motions.

The values of α under control condition, i.e. microtubules gliding in the absence of actin (α = 1.74 : 1.51 – 1.89 (median : 25^th^ quartile – 75^th^ quartile), *N* = 666, v = 9, *n* = 3) and in the presence of branched actin (α = 1.72 : 1.49 – 1.90, *N* = 479, v = 10, *n* = 3) were not different (Mann-Whitney U-test of equal medians, p = 0.71). In the linear actin, however, was the diffusion exponent (α = 1.55 : 1.21 – 1.84, *N* = 380, v = 10, *n* = 3) significantly reduced compared to control (p = 1.55 · 10^-11^). Increasing the motor surface density above a factor of 1.5 led to the diminishing of this effect (Fig. S2 b). Microtubules longer than 10 μm, which in each group represent approximately 25 % of microtubules (Fig. 2 e), tend to evince larger α (1.87, 1.84 and 1.75 (median) for the control condition, branched actin, and linear actin, respectively) compared to shorter microtubules (1.72, 1.68, and 1.46)(Fig. S2 c). Microtubules with diffusive and sub-diffusive trajectories (α ≤ 1) were almost three times more likely to be found in linear actin than in the other two groups (14 % vs 5 % of the corresponding populations), indicating increased confinement. Tethering the actin filaments partially to the substrate by biotin-neutravidin link (Methods) did not change this behaviour (Fig. 2 f, g), providing a piece of strong evidence that the variation in the strength of the actin depletion from the volume of the channel is not the cause of the observed difference. Bundling the filaments by α-actinin (a cross-linker that promotes the formation of actin bundles), in contrast, leads to a strong decrease in the diffusion exponent (Fig. 2 f, g), indicating that the architectural features of the actin network, which practically determine the interaction landscape experienced by microtubules, are the key parameter controlling the trajectories. We assume that the reduction of actin flexibility and the local augmentation of the interaction surface, both resulting from actin bundling, are responsible for the this observed decrease of the diffusion exponent

To link these results to the actin architecture, we first quantified the geometrical organisation of actin. Isotropic parameter analysis (Methods) confirmed that the branched network is highly anisotropic, unlike the linear actin. There is virtually no overlap of the isotropic parameter value between the two architectures regardless of the scale over which it is measured (Fig. 3 a, b). Nevertheless, the spread of the values of the isotropic parameter from several replicates led us to couple the microtubule diffusion exponents to the isotropic parameters of actin in which they were observed. A plot of these coupled values revealed a functional relation between the actin isotropy and the diffusion coefficient of microtubules gliding in this actin organisation (Fig. 3 c). Regardless of the actin architecture, the higher actin isotropy leads, on average, to a higher diffusion exponent of microtubules, following an empirical power-law increase.

**Figure 3.**
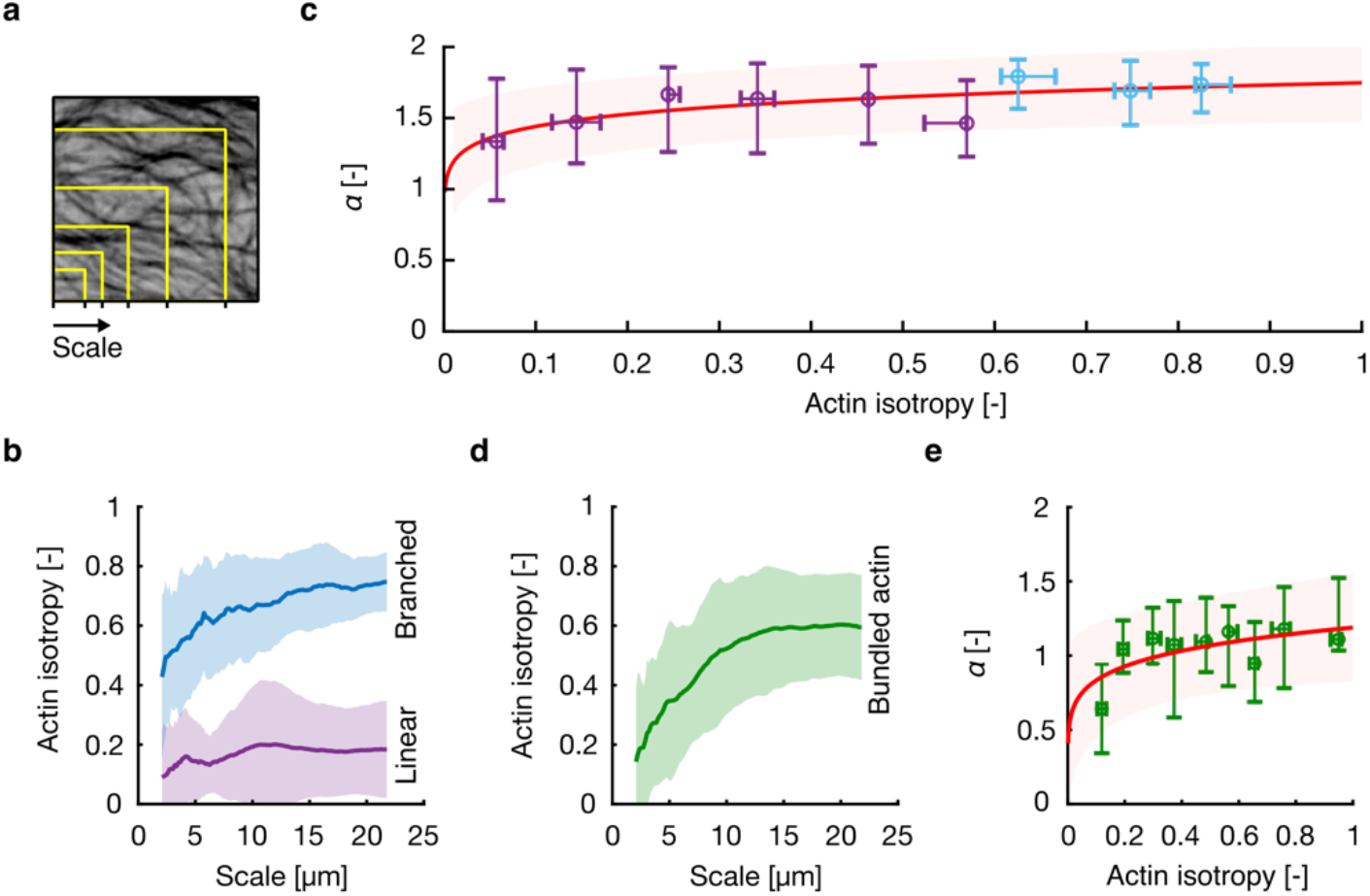
Microtubule motility depends on the isotropy of actin network. **a**, Illustration of the concept of scale. **b**, Actin isotropy as a function of scale for linear and branched actin. Data are represented as mean (solid line) ± s.d. (shadowed area). Branched actin: v = 10, *n* = 3. Linear actin: v = 10, *n* = 3). **c**, The diffusion exponent, a, as a function of actin isotropy, *L*. Data are binned and represented as median with whiskers extending to lower and upper quartiles. The red curve represents the power-law fit to medians within each bin (R^2^ = 0.6), and the shadowed area represents the 95% confidence interval. **d**, Actin isotropy as a function of scale for bundled actin. Data are represented as mean (solid line) ± s.d. (shadowed area). **e**, Diffusion exponent of microtubules gliding in bundled actin vs actin isotropy corresponding to the microtubule length. Central marks indicate median and the error bars indicate 75^th^ and 25^th^ percentiles of the data. The red curve represents the power-law fit (R^2^ = 0.44) to medians within each bin, and the shadowed area represents the 95% confidence interval.

We tested this observation further by repeating the same analysis in the linear actin network bundled by a-actinin. On a large uniform scale, contrary to our previous observations, we did not observe a clear functional relation (Fig. S3). However, we note that the bundled network has dramatically altered isotropy, which is strongly dependent on the scale because the mesh size of the network is relatively large (Fig. 3 d). Once we corrected the scale of the actin isotropy to the scale of each microtubule, practically adjusting the scale to the size of the microenvironment experienced by the microtubule, the clear functional pattern re-appeared (Fig. 3 e).

Having shown that microtubules sense and respond to actin architecture and organisation, we aimed to scrutinise the effect of the actin microenvironment on the gliding velocity of microtubules. Tracking the microtubules revealed that their mean gliding velocity has a rather wide distribution and is, on average, lower in the presence of actin (0.4 μm·s^-1^ (median) in both architectures) than at control condition without actin (0.5 μm·s^-1^) (Fig. 4 a), confirming the initial visual observations (Fig. 2 a). Increasing the concentration of ATP did not lead to an increase in the gliding velocity, demonstrating that the experiments were performed at the saturation condition (*33*) and confirming thus that the lower velocity in the presence of actin is not an experimental artefact stemming from inconsistencies in ATP concentration (Fig. 4 b). The difference between the best Michaelis-Menten velocity data fits (assuming that the Michaelis constant is not affected by the presence of actin) suggests that the interaction of microtubules with actin is asymptotically responsible for a 23% reduction of the gliding velocity. If the force-velocity curve of gliding microtubules were linear, this reduction would indicate that the friction force would, on average, reach 23% of the force required to stall the microtubule.

**Figure 4.**
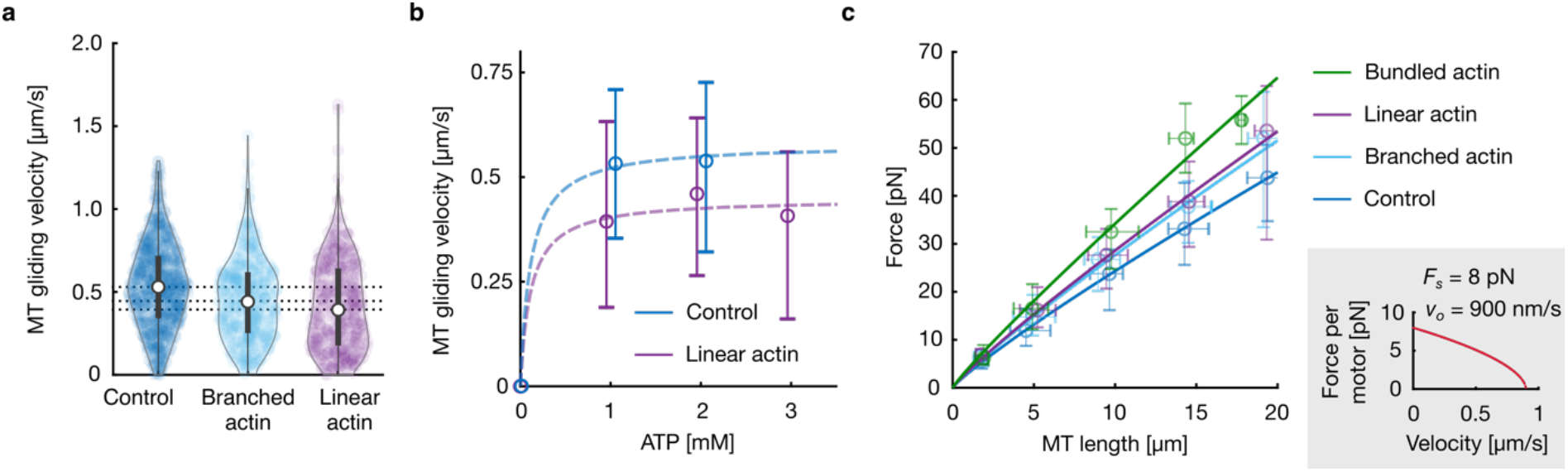
Analysis of the friction between actin filaments and microtubules. **a**, Violin plots of the velocities of microtubules gliding in the absence of actin and in the linear and branched actin networks. Individual data points are accompanied by violin plots: Central marks represent the median, top and bottom edges of the box indicate the 75th and 25th percentiles, respectively. Whiskers extend the 95% confidence intervals. **b**, Microtubule gliding velocity as a function of ATP concentration. Data are represented as median, and whiskers extend to lower and upper quartiles. The dashed line represents the Michaelis-Menten model. **c**, The force opposing microtubule movement as a function of microtubule length. Data are binned and represented as median with whiskers extending to lower and upper quartiles. Solid line plots are fits to the binned data. The force-velocity model used for the force estimation is shown in the inset.

The viscous drag, as well as the friction, are both functions of the microtubule length. As the samples of microtubules at the three conditions have similar yet not precisely identical lengths (Fig. 2 e), we next decoupled the distributions. We estimated the forces opposing microtubule gliding based on the three-parameter model of the force-velocity relation (*34, 35*) from the average gliding velocity, 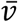, as 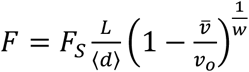, where *F_S_* is the stall force of the motor, *L* is the length of the microtubule, 〈*d*〉 is the average distance between microtubule-attached motors, *v_o_* is the velocity of the unloaded motor, and *w* is a scaling parameter (*35*). For the calculations, the following values of the parameters were used: *F_S_* = 8 pN, 〈*d*〉 = 1.8 μm, *v_o_* = 0.9 μm s^-1^, and *w* = 1.8 (*36*). In all groups, the opposing force increases with the microtubule length (Fig. 4 c). The control group exhibited a substantial, slightly non-linear trend with viscous drag from the medium and friction with the substrate as the assumed main contributing factors. We interpret the additive force in the presence of actin networks, which has a linear trend (R^2^ > 0.97 for all groups) since the majority of observed velocities fall within the linear part of the force-velocity curve, as the friction force between actin filaments and the microtubule. The addition is, per unit of microtubule length, 0.33 pN·μm^-1^ (95% confidence interval: 0.15 – 0.52 pN·μm^-1^) in branched and 0.43 pN·μm^-1^ (95% confidence interval: 0.25 – 0.60 pN·μm^-1^) in linear actin. We compacted linear actin with a-actinin to investigate this additive opposing force further. Doing so increased the opposing force by a factor of 2 per unit length of a microtubule compared to linear actin alone, providing evidence that the opposing forces are of frictional character.

## Discussion

By exploiting the well-documented sensitivity of motile microtubules in the gliding assay to external physical cues (*15, 19*), we studied the response of microtubules in this assay to the presence of actin filaments. Since the lengths of microtubules had a similar distribution in all the experimental groups reported here, and the length-decoupled data retained systematic differences, we can conclude that the observed differences between the groups were solely due to the actin condition. Within the groups, shorter microtubules, which are associated with fewer surface-bound kinesin-1 motors, were more sensitive to the presence of actin. The torsional compliance of kinesin motors provides enhanced rotational freedom to microtubules which are tethered to a very small number of motors (*37, 38*). External cues can, therefore, deflect the trajectory of the short microtubule more easily than that of long microtubules, where additional force is required to deform the microtubule lattice. Along the same lines, the motor surface density is an important parameter influencing the sensitivity of microtubules in the gliding assay. Lower motor density enables microtubule tips to explore a larger area by thermal fluctuations and, at the same time, it limits the ability of microtubules to overcome hindering forces, even though this relationship is not linear (*35*). Our observation that the increase of the motor density suppresses the effect of actin on the diffusion exponent of microtubules agrees well with this established understanding of the sensitivity of the gliding assay. We speculate that incorporating kinesin motors into the lipid bilayer, which regulates the force that motors can effectively produce by enabling their lateral diffusivity (*39–41*), could be instrumental for the further development of our assay. It would also allow for patterning both types of actin architectures simultaneously.

Our results show that the most dramatic effects of actin on microtubules, the guidance and frictional deceleration, were observed in anisotropic unbranched actin network, which enables tight alignment between the cytoskeletal polymers. This level of alignment could not be achieved in a branched actin network, where the microtubule is equally likely to encounter actin filament in any direction, making the mechanical environment and the interaction landscape highly isotropic. To observe some alignment, the distance between branching nodes would have to become comparable with the size of a microtubule or higher. Even though such a condition is experimentally inaccessible without changing the density of the actin network, the experiment with bundled actin network, which practically delivers a larger mesh size, confirms this view.

The absence of transport or visible deformation of actin filaments by motile microtubules indicates that microtubules if they become physically blocked by actin, cannot surpass the viscous drag of the medium on the actin network with which they interact. Such situations are predominantly transient as the microtubules can change their gliding direction. The intriguing, nevertheless, rare observation of quasi-oscillatory microtubule movement indicates elastic coupling (*42*) to actin by local strains in the actin mesh (*43*). In the future, direct force measurements shall help resolve this question in greater detail. Generally, the dominant forces hindering the microtubule movement are friction (*44*) and compaction (*45*). Friction has recently been established as an important factor in the behaviour of active materials (*46*). In our assay, the friction has various components, and we estimated the additive contribution caused by the presence of actin filaments. While the friction between cytoskeletal filaments is generally assumed non-linear (*44*), this friction in our assay has fairly linear character due to the limited width of the velocities population, which largely fall within the linear regime of the force-velocity model used (*34–36*). The compaction, which is caused by the presence of methylcellulose, is essential to i) offset the electrostatic repulsion between microtubules and actin filaments and ii) stabilise the actin mesh at the coverslip surface. Although we could not vary the concentration of methylcellulose in our experiments due to the use of untethered actin filaments, we can assume that the variation of the depletant concentration would modulate the strength of the interaction, similarly to other experiments with cytoskeletal polymers (*27*). Both friction and compaction depend on the geometry of the interacting surfaces and, thus, on the actin architecture in our experiments. The possibility of tight alignment between microtubules and linear actin architecture shall maximise the compaction, and frictional forces and our observations agree with this assumption.

The recognition of actin architecture by microtubules in mammalian cell cortex is governed by the microtubule dynamics (*47, 48*), crosslinking (*49, 50*) and retrograde actin flow (*51–53*), which, in contrast to our assay, result in negligible movement of microtubule lattice relative to actin network (*54, 55*). Nevertheless, our findings highlight the importance of the effects that non-specific steric interactions may have in biological or bio-inspired systems (*45, 56*).

While the interaction between linear actin filaments and microtubules in gliding assays has been studied recently (*29, 30*), here we demonstrate that the actin architecture regulates the motility of kinesin-propelled microtubules up to direct steering. We foresee a large potential for employing these findings in active materials and nano-devices, especially with patterned actin architectures, or in systems supporting dynamic switch of the actin architecture from linear to branched and *vice versa*. Since gliding assays have recently transcended beyond cytoskeletal biopolymers (*57*), the possibility of controlling the gliding properties by the anisotropy of the soft polymer layer becomes even more controllable and thus better applicable.

## Methods

### Protein purification

Bovine brain tubulin was isolated by recurrent polymerisation and depolymerisation (*58*) and purified from associated proteins (MAPs) by cation exchange chromatography (*59*). Rabbit skeletal muscle actin was purified from acetone powder and gel-filtered (*60*). Fluorescence labelling by Atto488 (tubulin) and Alexa568 (actin) fluorophores was accomplished using the NHS ester coupling. Recombinant, truncated kinesin-1-GFP motor was expressed in *E. coli* cells and purified as described previously (*61*). a-actinin was prepared by bacterial overexpression of pGex4T-1-ACTN4-6xHis construct and its consequent purification (*62*). Microtubules were polymerized from 4 mg/ml 1:4 mixture of labelled and unlabelled tubulin for 2 h at 37 °C in BRB80 (80 mM PIPES, 1 mM EGTA, 1 mM MgCl_2_, pH 6.9) supplemented with 1 mM MgCl_2_ and 1 mM GMPCPP. At the end of the polymerisation, microtubules were centrifuged for 30 min at 18000 x g. The pellet was resuspended in BRB80 supplemented with 10 μM taxol, and the microtubules were used within 5 days. Actin seeds for experiments with tethered actin were pre-polymerised from 5 μM, 2.3% biotinylated actin in the presence of 1.2 μM rhodamine-phalloidin.

### Motility assay

The imaging chamber was formed from NaOH-activated glass coverslips using double-sided tape. The surface of the channel was functionalised with GFP antibodies (Invitrogen, A-11122, 100 μg ml^−1^, 3 minutes, in 10 mM Hepes pH 7.2, 5 mM MgCl_2_, 1 mM EGTA, and 50 mM KCl), and the remaining available surface was passivated by BSA (Bovine serum albumin, 1% w/v, 5 minutes). Next, kinesin-1-GFP molecules (6 or 60 μg ml^−1^in wash buffer: 10 mM HEPES buffer (pH 7.2), 16 mM PIPES buffer (pH 6.8), 50 mM KCl, 5 mM MgCl_2_, 1 mM EGTA, 20 mM dithiothreitol (DTT), 3 mg ml^−1^ glucose, 20 μg ml^−1^ catalase, 100 μg ml^−1^ glucose oxidase, and 0.3% w/v BSA) were specifically attached to the antibodies (3 minutes), and the chamber was washed with three-times its volume of the wash buffer. Microtubules (1:300 dilution in wash buffer) were introduced in the channel and allowed to bind kinesin motors (1 minute). The channel was perfused by three times its volume of the wash buffer again to remove unattached microtubules, and, finally, the imaging buffer containing ATP was introduced (10 mM HEPES pH 7.2, 16 mM PIPES pH 6.8, 50 mM KCl, 5 mM MgCl_2_, 1 mM EGTA, 20 mM dithiothreitol (DTT), 3 mg ml^-1^ glucose, 20 μg ml^-1^ catalase, 100 μg ml^-1^ glucose oxidase, 2.67 mM ATP, 1 mM GTP, 0.3% w/v BSA, and 0.327% w/v methylcellulose (63 kDa, Sigma-Aldrich, M0387)). For experiments with actin, the imaging buffer was supplemented with 4 μM actin. Tethered actin filaments were prepared by elongation of pre-polymerised actin seeds attached to the coverslip via neutravidin by addition of 4 μM actin monomers. Bundled actin network was generated by supplementing 4 μM actin with 25 nM α-actinin. To generate a branched actin network, 80 nM Arp2/3 and 200 nM GST WA accompanied actin. The chamber was sealed by a capillary tube sealant (Vitrex) to prevent evaporation during the imaging. Surface motor density was evaluated by a photometric comparison of the fluorescence signal of the motors with the intensity of individual dimeric kinesin-1-GFP motors.

### Imaging

Total internal refraction fluorescence (TIRF) microscopy was used to image microtubules and actin filaments. The inverted microscope (Eclipse Ti, Nikon) was equipped with 100X 1.49 N.A. oil immersion objective (UApo N, Olympus), 491 nm and 561 nm lasers (Optical Insights) and an iLas^2^ dual laser illuminator (Roper Scientific) for the excitation of the Atto488 and Alexa568 fluorophores, respectively, Dual-View beam splitter (Optical Insights), and Evolve 512 EMCCD camera (Photometrics). Metamorph software (v. 7.7.5, Universal Imaging) was used to control the imaging. Images were taken every second.

### Image and data processing

Image and data processing were performed using Fiji (*63*) and MATLAB (MathWorks, Inc.). Dedicated Fiji plug-ins were used to generate maximum projections (Z project), colour-coded maximum projections (Temporal-Color Code), and kymographs (Multi Kymograph), and to monitor actin growth (Plot Z-axis profile). Microtubules were tracked using FIESTA (*63*), with the background of microtubule fluorescence images removed for this purpose. In the following analysis, we included only microtubules that were observable for at least 10 s and circumscribed trajectory of at least 6 μm to eliminate trajectories too short for the MSD curve fitting. The MSD curves and values of the diffusion exponent were obtained from the @msdnalyzer MATLAB class (*65*). Although some MSD curves suffered from offset arising from the localisation uncertainty, their slope used for analysis remains unaffected (*66, 67*). As a measure of actin isotropy, *L*, we used the circular variance of the director field of actin orientation (*68*). The value of *L* was estimated from the histogram of actin orientations obtained from the OrientationJ (*69*) plug-in for Fiji with the help of Matlab-FiJi interface MIJ (*70*) as:

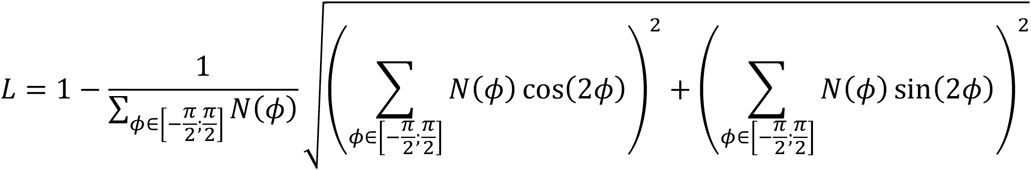

where *N*(*ϕ*) is the number of pixels oriented in the direction of angle *ϕ*. Statistical plots were generated in MATLAB using scripts by Bastian Bechtold for visualising violin plots. Statistical tests were performed using Statistics and Machine Learning Toolbox for MATLAB. The number of experiments, *n*, the minimum number of technical replicates in the experiment, v, and the number of data points, *N*, are indicated in the figures’ captions.

## Acknowledgement

We thank Laura Aradilla Zapata (Saarland University) for her critical comments on our results and Matthieu Gélin for inspiring this research. This work was supported by the European Research Council, Consolidator Grant 771599 (ICEBERG) to MT and Advanced Grant 741773 (AAA) to LB. OK was partially supported by Pôle emploi (7820342X). Our imaging platform is supported by the Laboratory of Excellence Grenoble Alliance for Integrated Structural & Cell Biology (LabEX GRAL)(ANR-10-LABX-49-01) and the University Grenoble Alpes graduate school (Ecoles Universitaires de Recherche, CBH-EUR-GS, ANR-17-EURE-0003).

## Author contribution

Conceptualization, LB, MT, OK; Methodology, OK, JG, CG; Investigation, OK; Formal Analysis, OK; Data curation, OK; Validation, OK; Resources, OK, JG, CG; Writing, OK; Visualization, OK; Supervision, MT, LB; Funding acquisition, MT, LB.

---

**Figure S1.**
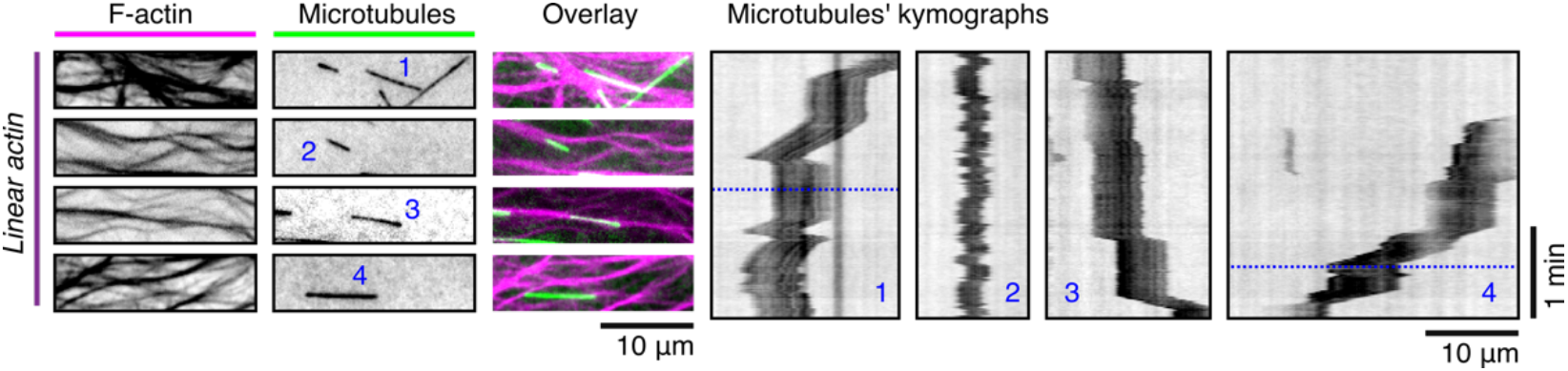
Complex motility of microtubules in linear actin. Intensity inverted fluorescence micrographs and corresponding kymographs of unusual motility of microtubules in linear actin. The horizontal blue line in the kymograph indicates the time when the corresponding fluorescence micrograph is shown if this time differs from 0.

**Figure S2.**
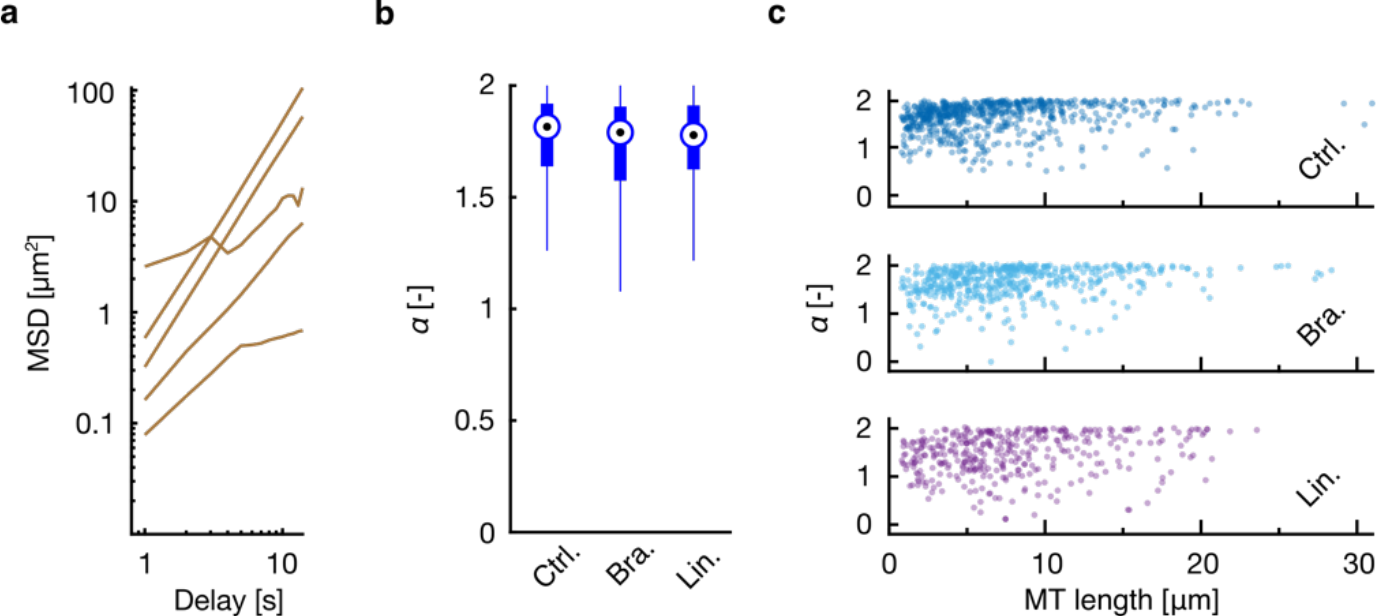
**a**, Mean squared displacement of microtubules from Fig. 2 b. **b**, The comparison of the diffusion exponent of microtubules gliding in the three types of linear actin at elevated motor surface density (Control: *N* = 287, v = 21, *n* = 2. Branched actin: *N* = 587, v = 15, *n* = 8. Linear actin: *N* = 352, v = 6, *n* = 2). **c**, Scatter plot of the diffusion exponent vs microtubule length.

**Figure S3.**
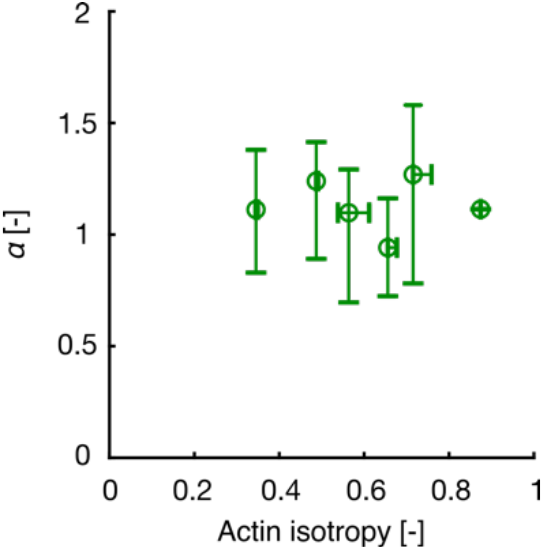
Diffusion exponent of microtubules gliding in bundled actin vs maximum scale actin isotropy (*N* = 114, v = 10, *n* = 3).

